# Monitoring hPSC genomic stability in the chromosome 20q region by ddPCR

**DOI:** 10.1101/2023.07.14.549021

**Authors:** Caroline Becker, Sema Aygar, Laurence Daheron

## Abstract

Copy number increases involving chromosome 20q with gain of the gene *BCL2L1* are a prevalent form of genomic instability in hPSC. In addition to large aneuploidies, findings in this region often include microamplifications that are too small to detect by G-banded karyotyping. Gene editing procedures warrant especially close monitoring of 20q genomic stability because they involve p53-activating stressors that select for the survival of *BCL2L1*-aneuploid cells. Here we describe an optimized strategy for detecting *BCL2L1* copy number increases in hPSC cultures using duplexed droplet digital PCR (ddPCR) with genomic DNA or cell lysate as the starting material. The procedure consists of droplet generation, thermocycling, droplet reading and data analysis. The expected result is a copy number estimate derived by comparing the number of droplets positive for *BCL2L1* to the number positive for a reference template, *PVRL2*. This procedure generates same-day screening results for 1 to 96 samples, providing a convenient option for screening hPSC cultures that is easily integrated into a gene editing workflow.

## Introduction

### Prevalence

Genomic integrity screening of human pluripotent stem cells (hPSCs) commonly reveals copy number increases affecting the q arm of chromosome 20. In addition to large karyotypic abnormalities such as trisomy 20 and isochromosome 20q, findings often include microamplifications in the 20q11.2 cytoband that are not reliably detected by G-banded karyotyping. A 2011 global screening effort identified gains at 20q11.2 in 25% of karyotypically normal hPSC lines [1], while a 2020 meta-analysis identified CNVs in this region as the most commonly reported recurrent genetic abnormality in hPSCs, accounting for 22.9% of reported abnormalities [2]. Here we briefly review the available information about 20q11.2 copy number increases in hPSCs and describe a step-by-step protocol for detecting these variations using droplet digital PCR (ddPCR).

### Mechanism and Risk Factors

The 20q11.2 region is adjacent to a repetitive sequence element, which likely creates a vulnerability to structural variations such as copy number amplifications [1]. Once the region is amplified in one cell, increased expression of the gene *BCL2L1* confers a selective advantage that promotes the expansion of the affected subpopulation within the culture. The main product of *BCL2L1* in hPSCs is the anti-apoptotic protein BCL-XL; accordingly, protection against apoptosis mediates the selective advantage associated with 20q11.2 amplification [3, 4].

BCL-XL functions as a brake on p53-dependent apoptosis by binding and sequestering cytoplasmic p53 [5]. Thus, any form of stress that activates the p53-dependent apoptotic pathway can promote the selective expansion of cells that overexpress BCL-XL. One such form of stress is the loss of cell-cell contact caused by single-cell dissociation and low-density plating [3, 6, 7]. Another is the induction of DNA double-strand breaks, including Cas9 nuclease cutting [8] and damage incidental to Cre-mediated recombination [9]. Gene editing workflows often involve exposure to multiple stressors, and indeed, BCL-XL overexpression has been found to promote hPSC survival during CRISPR-Cas9 gene editing [10]. Therefore, gene editing procedures warrant especially close monitoring of 20q11.2 genomic integrity.

### Impacts on Research

Existing work has demonstrated functional consequences of 20q11.2 amplification outside of survival advantage, including significant alterations to TGF-β- and SMAD-mediated signaling, and impairment of directed differentiation into neuroectoderm [11]. Impaired differentiation into hematopoietic precursors has also been reported [12]. In addition to the *BCL2L1* gene, amplifications of the region also result in the collateral gain of nearby genes, including the transcriptional regulator *ID1* and often *DNMT3B, a marker of undifferentiated hPSCs* [1]. Furthermore, *BCL2L1* copy number alterations have the potential to impact 3D tissue morphogenesis, since BCL-XL overexpression has been found in developmental contexts to promote the filling of lumens and ducts by migrating cells, while the absence of BCL-XL permits lumen clearance and cyst formation [13, 14]. In light of these findings, researchers should be aware that differentiation protocols developed using 20q11.2-aneuploid cells may require modifications, such as additional pro-survival factors, to achieve equivalent outcomes in euploid cells, and vice versa.

#### ddPCR Screening Method

ddPCR is one option in the toolkit for evaluating genomic integrity. The main benefits are the ability to obtain same-day screening results for 1 to 96 samples, and the ability to detect CNVs that are below the size limits of conventional karyotyping. The main drawbacks are the need for specialized equipment, and detection being limited to one or a few loci. Several groups have previously reported on the use of ddPCR for copy number screening in the 20q11.2 region [2, 15]. The general procedure consists of droplet generation, thermocycling, droplet reading and data analysis, and the result is a copy number estimate derived by comparing the number of droplets positive for *BCL2L1* to the number of droplets positive for a reference template. In this protocol, we provide detailed advice for implementing ddPCR screening based on observations from over 900 assay runs and offer optimization strategies addressing several practical challenges.

#### Selection of Reference Assay

One issue impacting ddPCR-based CNV analysis in hPSCs is the tendency of assays to exhibit a bias in which diploid samples (including validated controls) read either slightly above, or slightly below, 2.0 copies of the target, depending on the assay. We encountered this phenomenon during protocol development, and examples can also be observed in previously published data [15]. When such mid-integer CNV calls cannot be traced to sample quality or mosaicism, they may be due to replication timing, i.e. how early or late in S-phase a given genomic region undergoes DNA replication in a given cell type [16]. If the target gene replicates earlier than the reference gene, then without cell synchronization, there will be a percentage of cells in the culture that have replicated the target locus but not the reference locus, resulting in an overestimate of the copy number. We note that since pluripotent cells spend more of their time in S-phase than other cell types, owing to a short G1 phase [17], bias due to replication timing would tend to be magnified in hPSCs. We therefore conducted a replication timing-aware search for the reference locus most suitable for CNV analysis of *BCL2L1*.

Although *PVRL2* (a.k.a. *NECTIN2*) is not a housekeeping gene, we recommend it as a reference gene for *BCL2L1* for several reasons. *PVRL2* is located on chromosome 19q, where copy number variations in hPSCs are rare, and a meta-analysis of published data did not identify any gains or losses at the *PVRL2* locus [15]. Furthermore, *PVRL2* has similar replication timing to *BCL2L1* in hPSCs, in contrast to reference loci used in other protocol versions such as *RPP30* [2] and *TERT* [15] (Fig. 1).

**Figure 1:**
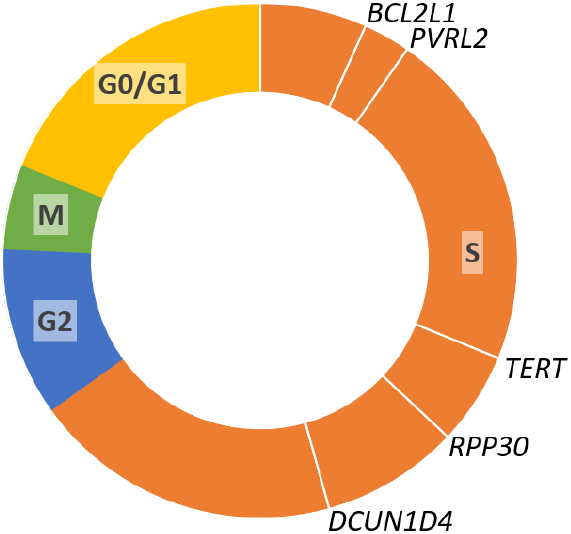
Cell cycle distribution of proliferating H1 and H9 human ES cells [17] with average replication timing of *BCL2L1* and potential reference loci in hPSCs. Replication timing based on Repli-chip from ENCODE/FSU, accessed via UCSC Table Browser [18-20]; cell lines H1, H7, H9, BG02ES, hFib2 iPS4 and hFib2 iPS5.

Finally, a *PVRL2* copy number assay is commercially available from Bio-Rad and performed well in both the manufacturer’s validation (exhibiting 95% specificity) and in our experiments.

To validate the assay design, we performed ddPCR screening on 23 hPSC genomic DNA samples that we previously screened by SNP array (Illumina Global Screening Array v1.0). 4/4 aneuploid control samples, each with 3-4 copies of *BCL2L1* based on SNP array, exhibited ddPCR measurements of 2.95-3.88 copies. 18/19 euploid control samples, each with 2 copies of *BCL2L1* based on SNP array, had ddPCR measurements of 1.89-2.15 copies, while the remaining sample had a ddPCR measurement of 2.28 copies.

#### Cell lysate option

While isolated genomic DNA offers the best precision, DNA extraction can be impractical in the case of screening large numbers of clones. To address this challenge, we offer a protocol modification for the use of cell lysate as the starting material. In our laboratory, a normal cell lysate result predicted a normal genomic DNA result 4-6 passages later in 98% of clones (N=40). Without pre-screening, 48% of clones were euploid at the *BCL2L1* locus after expansion (N=83).

### Application in hPSC Quality Control

This procedure is applicable as part of a comprehensive hPSC quality control strategy and can be integrated into gene editing workflows. To streamline the commitment of resources, we recommend routine screening of hPSC cultures at several points throughout the process of gene editing:

#### 1. Parental cell lines

Screening parental cells prior to editing allows the user to avoid targeting cell lines or stocks that already carry a *BCL2L1* copy number increase.

#### 2. Polyclonal pools

Screening after transfection, but before single cell cloning, allows the user to assess the impact of the editing step on the average *BCL2L1* copy number. If multiple conditions are tested, screening the polyclonal pools makes it possible to choose the pool with the best combination of editing efficiency and 20q11.21 genomic integrity.

#### 3. Picked clones

If the editing project includes single-cell cloning, clones with desired genotypes can be screened using a small portion of the sample material harvested for genotyping.

#### 4. Final product

After clones are chosen and expanded, a final screening step can help assure quality for downstream applications.

## Workflow

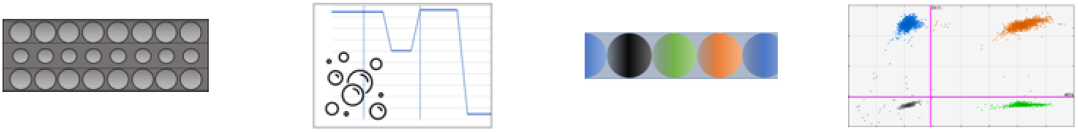

## Materials and Preparation

### Droplet Generation

#### Reagents

- ddPCR™ Supermix for Probes (No dUTP) (Cat. no. 1863023, Bio-Rad Laboratories, Hercules, CA, USA)
- PrimePCR™ ddPCR™ Copy Number Assay: BCL2L1, Human, FAM (Unique Assay ID: dHsaCP1000156, Bio-Rad Laboratories, Hercules, CA, USA)
- PrimePCR™ ddPCR™ Copy Number Assay: PVRL2, Human, HEX (Unique Assay ID: dHsaCP2506625, Bio-Rad Laboratories, Hercules, CA, USA)
- Restriction Enzyme HaeIII, 10,000 units/ml concentration (Cat. no. R0108S, New England Biolabs, Ipswich, MA)
- Nuclease-free water

#### Cell Lysis Reagents (Optional)

- UltraPure™ 0.5M EDTA, pH 8.0 (Cat. No 15575020, Thermo Fisher Scientific, Waltham, MA, USA)
- UltraPure™ 1M Tris-HCI, pH 8.0 (Cat. No 15568025, Thermo Fisher Scientific, Waltham, MA, USA)
- Triton™ X-100 solution, ∼10% in H2O (Cat. No. 93443, MilliporeSigma, Darmstadt, Germany)
- Proteinase K Solution (20 mg/mL), RNA grade (Cat. No 25530049, Thermo Fisher Scientific, Waltham, MA, USA)
- DPBS, no calcium, no magnesium (Cat. No 14190250, Thermo Fisher Scientific, Waltham, MA, USA)

#### Instruments

- QX200™ Droplet Generator (Cat. no. 1864002, Bio-Rad Laboratories, Hercules, CA, USA)
  - *Includes Droplet Generator Cartridge Holder (Cat. no. 1863051)*

#### Consumables

- 8-strip PCR tubes
- DG8™ Cartridges for QX200™/QX100™ Droplet Generator (Cat. no. 1864008, Bio-Rad Laboratories, Hercules, CA, USA)
- DG8™ Gaskets for QX200™/QX100™ Droplet Generator (Cat. no. 1863009, Bio-Rad Laboratories, Hercules, CA, USA)
- Droplet Generation Oil for Probes (Cat. no. 1863005, Bio-Rad Laboratories, Hercules, CA, USA)
- ddPCR™ Buffer Control for Probes (Cat. no. 1863052, Bio-Rad Laboratories, Hercules, CA, USA)

*Optional*

### Droplet Transfer, Plate Sealing and Thermocycling

#### Instruments

- PX1_**TM**_ PCR plate sealer (Cat. no. 181-4000, Bio-Rad Laboratories, Hercules, CA, USA)
- Thermocycler with 96–deep well reaction module

#### Consumables

- ddPCR™ 96-Well Plates (Cat. no. 12001925, Bio-Rad Laboratories, Hercules, CA, USA)
- PCR Plate Heat Seal, foil, pierceable (Cat. no. 1814040, Bio-Rad Laboratories, Hercules, CA, USA)

### Droplet Reading

#### Instruments

- QX200™ Droplet Reader (Cat. no. 1864003, Bio-Rad Laboratories, Hercules, CA, USA) *This protocol has not been tested with the QX100 system, but we expect the procedure and results to be the same.*
  - *Includes QuantaSoft*_***TM***_ *Software (Version 1*.*7*.

##### Consumables

- ddPCR™ Droplet Reader Oil (Cat. no. 1863004, Bio-Rad Laboratories, Hercules, CA, USA)

## Procedure

### Optional: Preparation of cell lysate from 96-well plate

#### Timing: 90 min

Preparation of the 4X lysis buffer. First, prepare the following stocks in nuclease-free water and store at room temperature:

| Component | Final Concentration | Amount | Nuclease-free water |
| --- | --- | --- | --- |
| 0.5M EDTA, pH 8.0 | 12 mM | 240 μL | 9.76 mL |
| 1M Tris-HCl, pH 8.0 | 30 mM | 300 μL | 9.7 mL |
| Triton™ X-100 solution,<br>~10% in H<sub>2</sub>O | 6% | 6 mL | 4 mL |

Prepare 4X lysis buffer for one 96-well plate:

Prepare 4X lysis buffer for one 96-well plate:
| Component | Final Concentration | Amount |
| --- | --- | --- |
| 12 mM EDTA | 4 mM | 400 μL |
| 30 mM Tris-HCl | 10 mM | 400 μL |
| Triton X-100, 6% | 2% | 400 μL |
| Proteinase K (20mg/mL) *freshly added* | 1% | 12 μL |

Pick clones into a 96-well plate and culture for 7-10 days.

During the first passage, reserve 1/3 of each clone (approx. 20k cells) for lysis. Transfer the portion of cells for lysis to a 96-well PCR plate and add DPBS to a volume of 30 µL per well.

Add 10 µL of 4X lysis buffer to each well.

Heat the plate at 60°C for 60 min. followed by 95°C for 10 min.

Add 60 µL nuclease-free water to each well.

A portion of the lysate may be used for clone genotyping. Store the remaining lysate at -20°C. Once clones with desired genotypes are identified, proceed with ddPCR screening for those clones.

### Preparation of Reaction Mixture

#### Timing: 10 min. + 10 min. per eight samples

Isolate genomic DNA or prepare cell lysate from each clone or cell line of interest.Using 8-trip PCR tubes, load each tube with the recommended amount of DNA according to the chart below. **Cell Lysate Option**: Prior to pipetting, centrifuge the lysate at full speed (∼18,000 x *g*) for 5 minutes to pellet debris and draw the sample from the top supernatant.

| Sample Type | Quantification Method | Amount per 25 μL reaction mixture |
| --- | --- | --- |
| Genomic DNA | Fluorescence | 150 ng dsDNA |
| Genomic DNA | Spectrophotometry | 350 ng nucleic acid |
| Cell Lysate | N/A | 4 μL |

Add nuclease-free water to a final volume of 10 μL.

*Note: For quantification of genomic DNA by spectrophotometry, use RNase A treatment during DNA extraction*.

*Note: Samples may be run with or without technical replicates. A single well per sample is sufficient to calculate a CNV estimate and Poisson confidence intervals reflecting theoretical technical error. By adding replicates, the user may also calculate total error*.

*Note: Droplet generation requires multiples of eight samples. If not running a multiple of eight DNA samples, you will need to fill the remaining DG8 cartridge wells with an appropriate solution: use either ddPCR*^***TM***^ *Buffer Control for Probes diluted with an equal volume of water or prepare additional reaction mixture with water in place of DNA*.

Prepare a master mix according to the chart below. Prepare enough master mix for your number of samples, plus 10% to account for pipetting loss.

| Component | Amount per sample | Final Concentration |
| --- | --- | --- |
| Supermix for Probes (No dUTP) | 12.5 µL | 1X |
| Copy Number Assay: BCL2L1, FAM | 1.25 µL | 900 nM each primer/<br>250 nM probe |
| Copy Number Assay: PVRL2, HEX | 1.25 µL |  |
| Restriction Enzyme HaeIII (undiluted) | 0.25 µL | 0.1 unit/µL |

Add 15.25 µL of the master mix to each 10 µL sample and mix well. While setting up the next step, leave the reaction mixtures at room temperature for 10 minutes to allow the restriction digest to run to completion.

*Note: 20 µL of each 25*.*25 µL reaction mixture will be used for droplet generation. The excess volume is used to ensure that there is no shortfall when loading the droplet generation cartridge*.

### Droplet Generation

#### Timing: 5 min. per 8 samples

*Note: Instructions for the following two sections are adapted from the Bio-Rad QX200 Droplet Generator Instruction Manual. Please refer to this manual for detailed safety, regulatory and maintenance information*.

Open the cartridge holder by gently applying pressure to the top and bottom hinges. Position a DG8 cartridge in the cartridge holder with the wells facing up and the notched edge oriented to the left. Close the cartridge holder by gently pushing the left and right sides together.

Using a single- or multi-channel pipette, load 20 µL of reaction mixture or 1X buffer control into the middle row of wells. Avoid creating bubbles; pipette into the bottom of the well at a slight angle and depress the pipette plunger only to the first stop.

Inspect each well carefully for bubbles. Remove any bubbles manually using a clean 10 µL pipette tip.

Using a single- or multi-channel pipette, load 70 µL of droplet generation oil into each well in the bottom row.

Attach a gasket across the cartridge by fitting the four hooks on the cartridge holder through the four holes in the gasket.

Press the button on the droplet generator to open the compartment. Place the prepared cartridge holder in the compartment and press the button again to begin droplet generation. The droplets are ready when the rightmost indicator light shows solid green. Timing: 2 min. per 8 samples

Retrieve the cartridge holder from the droplet generator.

Discard the gasket but do not open the cartridge holder. Check that each well of the top row contains a cloudy mixture, indicating the presence of droplets.

### Droplet Transfer and Plate Sealing

#### Timing: 5-10 min

*Critical Step: Once droplets are formed, work carefully but promptly. Do not pipet up and down or otherwise disturb the droplets more than necessary*.

Using a pipette set to 40 µL (a reliable multi-channel pipette is preferred if available), place the tips into the bottoms of the droplet-containing wells at a 45-degree angle. Slowly collect the well contents and transfer them to the ddPCR 96-well plate by slowly dispensing down the side of the well.

If additional droplets remain in the cartridge, add them to the same well of the 96-well plate by repeating the same procedure. Discard the cartridge.

Repeat droplet generation and transfer until all samples have been processed.

Remove the plate support block (if present) from the plate sealer and set it aside. Set the temperature to 180°C and time to 5 seconds.

When the temperature reading has reached 180°C, open the plate sealer and insert the support block with the 96-well side facing up. Place the 96-well plate containing droplets in the support block. Place a foil seal on top of the plate with the red stripe facing up. Press seal.

*Pause Point: The sealed 96-well plate may be stored at 4°C for up to four hours*.

### Thermocycling

#### Timing: 1 hour 45 min

Place the sealed 96-well plate in the deep-well thermocycler and run PCR according to the following program:

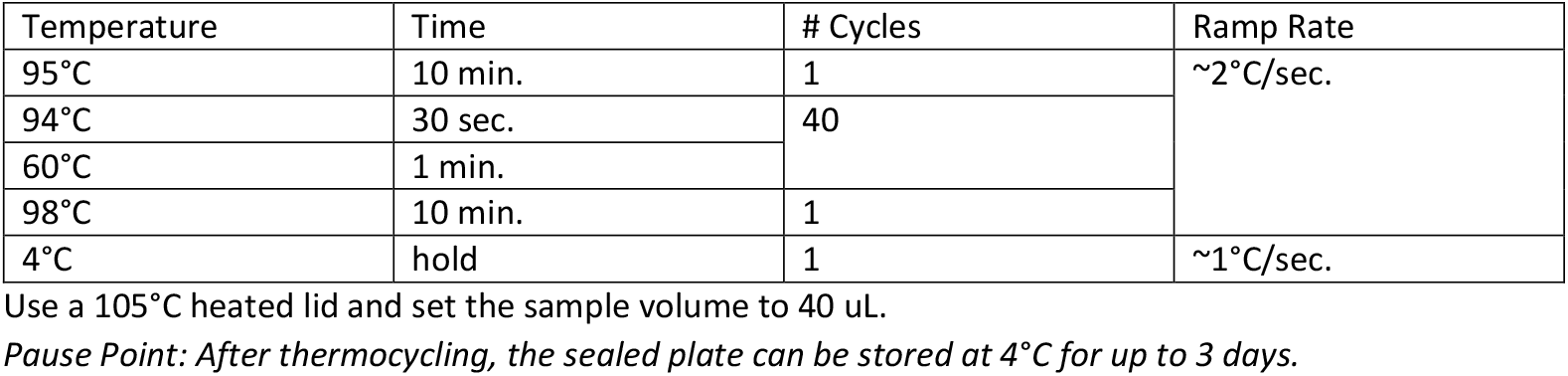

### Droplet Reading

*Instructions for the following two sections adapted from Bio-Rad QX200 Droplet Reader and QuantaSoft Software Instruction Manual. Please refer to the manual for detailed safety, regulatory and maintenance information*.

#### Setup time: 5-10 min

##### Droplet reader run time: 4 min. + 1.25 min. per sample

Open the plate holder by lifting the left and right clips. Seat the sealed 96-well plate in the base of the plate holder and place the lid on top. Push down on the clips to lock the 96-well plate in place.

Press the button on the front of the droplet reader to open the compartment. Seat the plate holder containing the 96-well plate in the compartment and close the compartment.

Launch QuantaSoft on the connected computer.

To set up a new assay, hold down the shift key and click to highlight the sample-containing wells of your 96-well plate on the plate diagram.

Still holding the shift key, double-click any of the highlighted wells to access the well settings.

Under Sample, select Experiment: CNV2 and Supermix: ddPCR Supermix for Probes (no dUTP). Under Target 1, type BCL2L1 and select Type: Ch1 Unknown.

Under Target 2, type PVRL2 and select type: Ch 2 Reference. Click Apply.

Next, double-click each sample-containing well individually and under Sample, type the sample name and passage number, then click Apply. Finally, click OK.

*Note: If you are running technical replicates and would like to enable merged data analysis, give the same name to all the replicates*.

In the left sidebar, click run, then click yes in the dialog box. Name and save your template file. After the run, the data will be saved in a plate file of the same name.

In the next dialog box, if your samples are ordered by columns, click “acquire data by columns.” Under dye set, select FAM/HEX. Click OK. Droplet reader run time is approximately 4 min. + 1.25 min. per sample.

## Data analysis

### Timing: 5 min

In QuantaSoft, click Analyze in the left sidebar.

For each sample, in the 96-well plate diagram in the upper-right panel, click the corresponding well. In the bottom panel, select the 2D amplitude tab. Examine the scatterplot to confirm that the droplet reader detected four populations.

Click on the scatterplot to place the quadrant gate such that one population is in each quadrant.

*Note: A small number of droplets falling in between the major populations is normal. Gate placement with respect to these droplets has a negligible impact on the assay result*.

After gating, refer to the table in the upper-left panel. The CNV statistic indicates the *BCL2L1* copy number estimate. The PoissonCNVMin and PoissonCNVMax columns reflect the Poisson 95% confidence interval for the CNV statistic.

Suggested ranges for interpretation are below, based on a panel of 19 hPSC control samples showing normal BCL2L1 and PVRL2 copy numbers by SNP array. The p-value column reflects the proportion of 2-copy samples expected to show an equal or higher CNV estimate based on the control dataset.

| CNV (gDNA method) | CNV (lysate method) | Interpretation | p-value |
| --- | --- | --- | --- |
| ≤2.15 | ≤2.47 | Normal |  |
| 2.16-2.18 | 2.48-2.51 | Borderline | $p \leq 0.1$ |
| 2.19-2.99 | 2.52-3.44 | Mosaic (mixed population) | $p \leq 0.05$ |
| 3.00 | ≥3.45 | Abnormal | $p < 0.00001$ |

*Note: The CNV (lysate method) column adjusts for an observed 15% overestimate of the BCL2L1 copy number using the method described under “Preparation of cell lysate from 96-well plate.” For samples prepared using methods that differ from this protocol, interpret the CNV statistic based on validated controls.*

## Timing

For eight gDNA samples the procedure takes approximately 2 hours and 10 minutes, consisting of:

- 30 minutes for droplet generation
- 80 minutes for thermocycling
- 15 minutes for droplet reading
- 5 minutes for data analysis

Each additional eight samples, up to 96, adds approximately 20 minutes to the total time.

## Expected Results

To evaluate whether the procedure provides consistent results under real usage conditions, we reviewed data from all hPSC samples that were run multiple times in separate experiments (samples A-I; 44 assays total). One sample (D) was a cell lysate; the rest were genomic DNA samples. Between-run variation impacted the result interpretation in only one case (sample F, outlier). QC data suggest that low DNA input (<40 ng) accounted for the largest errors (samples F and G, minima).

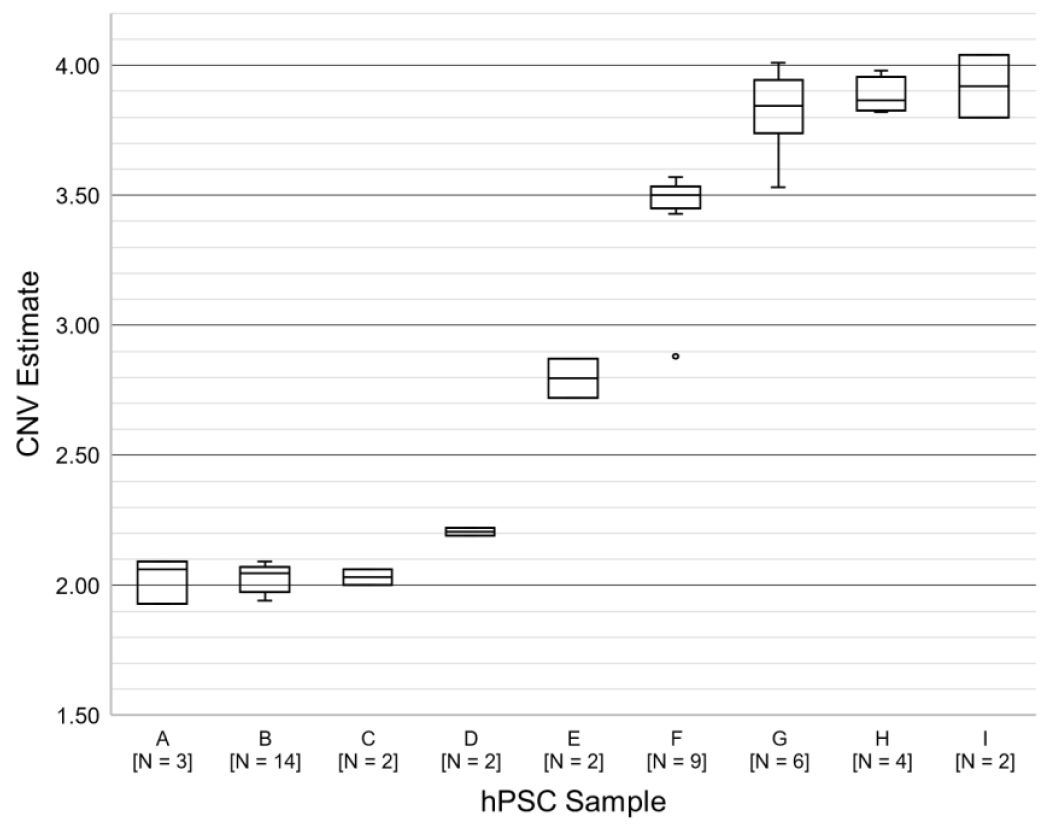

We retrospectively analyzed ddPCR data for 11 rounds of CRISPR/Cas9 targeting in hPSC (2 knock-out, 9 knock-in) including a total of 92 clones. All transfections in this analysis were performed on parental cell lines with a normal BCL2L1 copy number by ddPCR. For each targeting, different transfection conditions were tested such as electroporation voltage and ratio of cells to ribonucleoprotein. Polyclonal pools were screened by ddPCR after recovery from transfection. After subcloning one or more pools, clones with desired genotypes were screened for BCL2L1 copy number.

We obtained at least one BCL2L1-normal polyclonal pool in 9 of the 11 transfections. The other two pools were mosaic (mixed population) for gain of BCL2L1. Clones derived from normal pools had a normal BCL2L1 copy number 4.8x more often than the clones derived from mosaic pools: 62% (43/69) of clones from normal pools were normal, while only 3/23 (13%) of clones from mosaic pools were normal. These findings demonstrate the benefit of screening polyclonal pools for BCL2L1 CNVs prior to subcloning, to choose the edited pool with the best chance of yielding primarily euploid clones.

### Example for one gene editing project

We introduced the mutation c.397C>T (p.Arg133Cys) in the Notch3 gene in PGP-1 cell line. The parental line, PGP1 iPS at p19, had a BCL2L1 copy number of 2.23. To introduce the RNP and ssODN, the cells were transfected using the Neon system at three different voltages: 1,000v, 1,100v and 1,200v. The 3 pools of cells were tested for BCL2L1 copies. The pool of cells transfected at 1,000v had a copy number of 2.32, the pool at 1,100v was at 2.24 and the pool at 1,200v was 2.79. Although the targeting efficiency was higher at 1,200v, we selected the pool at 1,100v for single cell cloning to increase our change to obtain normal clones. After screening a 96 well plate, 11 clones were expanded (5 homozygous, 1 heterozygous and 5 untargeted control) and tested for BCL2L1. 6 out of 11 of these clones had a normal BCL2L1 copy number. The remaining clones had ∼3 copies.

## Troubleshooting

| Step | Problem | Possible Reason | Solution |
| --- | --- | --- | --- |
| Droplet generation | The sample is completely clear (no cloudy layer) following droplet generation | Bubbles or contaminants (e.g. excessive detergent) in the sample prevent droplet formation. | Centrifuge the sample at full speed (~18,000 x g) for 5 minutes prior to preparing the reaction mixture and draw the sample from the top supernatant. If unsuccessful, re-extract or purify the DNA sample.<br>Inspect the sample well and remove any bubbles prior to droplet generation. |
| Data analysis | <u>The droplet reader detected eight populations</u><br>Diagonal, smeared appearance of populations | Excessive pipetting force during the transfer of the droplets to the PCR plate can cause a subset of the droplets to coalesce or shear, causing the formation of a secondary cluster with higher signal amplitudes to appear next to each droplet population. | Gate each secondary cluster together with the larger cluster below and to the left of it. The data may still be interpreted normally, but with some loss of precision. |
|  |  |  | To reduce the likelihood of droplets coalescing, follow the instructions for droplet transfer carefully. |
| Data Analysis | <u>The droplet reader detected only two populations</u> | The same fluorescent label was used for both the target and reference gene. | Verify that the reaction mixture contains one FAM-labelled assay and one HEX-labelled assay. |
| Data Analysis | <u>Large error bars</u> | If the Poisson confidence interval for the CNV statistic (PoissonCNVMax-PoissonCNVMin) is wider than 0.25 copies, this indicates that either too much or too little DNA was present in the reaction mixture. In the droplet reader data, consult the column "CopiesPer20ulWell" for the PVRL2 reference assay; the preferred range is 20,000-40,000 copies. When using the cell lysate option, larger error bars are expected; increasing the amount of lysate is not recommended because a high concentration of lysis buffer interferes with droplet generation. | In future runs, adjust the amount of DNA input until the droplet reader reports between 20,000-40,000 copies of PVRL2 per 20 uL well<br>If cell lysate was used as the starting material, larger error bars are expected; increasing the amount of lysate is not recommended because carryover of the lysis buffer interferes with droplet generation. |
| Data Analysis | <u>Lack of negative (-/-) droplets</u> | Too much DNA was present in the reaction mixture. | In the droplet reader data, consult the column "CopiesPer20ulWell." In future runs, adjust the amount of DNA input until the droplet reader reports between |
|  |  |  | 20,000-40,000 copies of PVRL2 per 20 uL well. |

## Acknowledgements

We are grateful to the members of the Harvard iPS core facility for their helpful discussions

